# Physiological networks governing salinity tolerance potentials in *Gossypium hirsutum* germplasm

**DOI:** 10.1101/2019.12.16.877787

**Authors:** Kevin R. Cushman, Isaiah C. M. Pabuayon, Lori L. Hinze, Megan E. Sweeney, Benildo G. de los Reyes

## Abstract

Toxic ions begin to accumulate in tissues of salt-stressed plants after the initial osmotic shock. In glycophytes, the ability to mobilize or sequester excess ions define tolerance mechanisms. Mobilization and sequestration of excess Na^+^ involves three transport mechanisms facilitated by the plasma membrane H^+^/Na^+^ antiporter (SOS1), vacuolar H^+^/Na^+^ antiporter (NHX1), and Na^+^/K^+^ transporter in vascular tissues (HKT1). While the cultivated *Gossypium hirsutum* (upland cotton) is significantly more tolerant to salinity relative to other crops, the critical factors contributing to the observed variation for tolerance potential across the germplasm has not been fully scrutinized. In this study, the spatio-temporal patterns of Na^+^ accumulation at different severities of salt stress were investigated across a minimal comparative panel representing the spectrum of genetic diversity across the improved cotton germplasm. The goal was to define the importance of integrative or network effects relative to the direct effects of Na^+^ homeostasis mechanisms mediated by *GhHKT1, GhSOS1,* and *GhNHX1.* Multi-dimensional physio-morphometric attributes were investigated in univariate and multivariate statistical contexts, as well as the relationship between variables using structural equation modeling. Results showed that mobilized or sequestered Na^+^ may contribute to the baseline salinity tolerance, but the observed variance in overall tolerance potential across a meaningful subset of the germplasm were more significantly associated to antioxidant capacity, maintenance of stomatal conductance, chlorophyll content, and divalent cations, and other physiological interactions occurring through complex networks.

**One-Sentence Summary:** Variation in salinity tolerance potential across the tetraploid cultivated *Gossypium* germplasm is better explained by complex physiological networks rather than just cellular Na^+^ homeostasis.

## INTRODUCTION

High salt concentration in the soil impedes the ability of plant roots to extract water causing dehydration and osmotic stresses, with negative impacts to cellular processes that support vegetative and reproductive growth. With prolonged exposure, salt concentration within the plant could build-up leading to cellular toxicity. Injuries are due to physiological perturbations brought about largely by cell wall and membrane damage, ionic toxicity, and impairment of photosynthesis (Ashraf and Harris, 2004). Adaptive mechanisms for osmotic adjustment, tissue tolerance to accumulated Na^+^, and exclusion of excessive Na^+^ are evolutionarily conserved regardless of the plant’s inherent capacity for tolerance or avoidance (Nakayama et al., 2005).

Tolerance to osmotic stress is controlled by genes involved in long distance signaling (*SOS3*, *SnRKs*), osmotic adjustment (*P5CS*, *HKT1*, *SOS1*), and stomatal regulation (*ERA1*, *PP2C*, *AAPK*, *PKS3*) (Berthomieu et al., 2003; Ashraf and Harris, 2004; Nakayama et al., 2005; Munns and Tester, 2008). In concert, these processes slow down the rate of cell expansion in roots and young leaves (Yeo et al., 1991). Tissue tolerance involves efficient intracellular or intercellular compartmentalization of Na^+^ and Cl^-^ to prevent toxic effects in the cytoplasm through the sequestration of Na^+^ into the leaf vacuole by NHX1 and AVP proteins, and exclusion of Na^+^ by HKT1 and SOS1 proteins (Halfter et al., 2000). Exclusion and transport of Na^+^ prevent the rapid build-up of toxic concentrations in the leaves, through the regulation of net ion transport to the shoot by SOS3 and SnRK, avoidance of toxicity in the chloroplast mediated by PP2C and ERA1, reduction of long distance Na^+^ transport facilitated by HKT1 and SOS1, and by efficient sequestration of Na^+^ into the root vacuoles through NHX1 and AVP mediated mechanisms (Cheeseman, 1988; Apse et al., 1999; Blumwald et al., 2000; Halfter et al., 2000; Maser et al., 2002; Chinnusamy et al., 2004; Haro et al., 2005; Brini et al., 2007).

Numerous studies in Arabidopsis and several crop plants have shown that positive net gains in salinity tolerance can be achieved with the overexpression of critical genes involved in Na^+^ transport and sequestration. For instance, overexpression of *HKT1* has been shown to regulate the vertical distribution of Na^+^ and K^+^, and export of Na^+^ out of the xylem (Hauser and Horie, 2010). Overexpression of *HKT1* in the roots and lower leaves has also been shown to reduce Na^+^ concentration in the xylem sap, protecting the younger and more sensitive apical shoot meristems from toxic effects (Sunarpi et al., 2005; Davenport et al., 2007). Additionally, *HKT1* has an important role in phloem loading by regulating the removal or recirculation Na^+^ back to the roots (Berthomieu et al., 2003). The plasma membrane-associated SOS1 exports Na^+^ to the apoplast and intercellular spaces. It facilitates the removal of Na^+^ in root epithelial cells in association with pores and excretory glands (Shi et al., 2002; Ji et al., 2013). NHX1 sequesters Na^+^ into the vacuole to compartmentalize but not eliminate excess intracellular Na^+^ (Apse et al., 1999; Zhang and Shi, 2013).

Relative to most other crop plants, the tetraploid cultivated cotton (*Gossypium hirsutum* L.) has considerably better tolerance to dehydration as well as osmotic and ionic stresses, hence commonly grown in semi-arid or salinity-prone environments (Maas and Hoffman, 1977). Because domestication and subsequent cultivar development has been driven primarily by selection for seed yield and fiber-related attributes, it has always been perceived that limited variability exists across landraces and modern cultivars with respect to physiological traits that enhance the potential for osmotic and/or ionic stress tolerance. Despite this general assumption, plant breeders and physiologists have been in search for elite cultivars or landraces exhibiting relative superiority in terms of stress physiological attributes that can be combined with fiber yield and quality traits. A major hurdle is the lack of consensus set of physiological and biochemical parameters that can reveal meaningful variation across the germplasm that are also indicative of differences in overall potential. Nevertheless, efforts to screen the germplasm for variation for salinity tolerance at the whole plant level using both laboratory and field-based assays have revealed quantifiable inter-genotypic differences (Leidi and Saiz, 1997; Basal et al., 2006; Hemphill et al., 2006).

The *Gossypium* Diversity Reference Set (GDRS) is a germplasm panel representing the spectrum of geographic distribution as well as morphological and allelic diversity across landraces and cultivars collected worldwide (Hinze et al., 2016; Hinze et al., 2017). To uncover meaningful variation for salinity tolerance potential across the germplasm, we conducted an extensive study on a representative subset of haplotypes (*i.e., core-GDRS*) as minimal comparative panel for physiological attributes relevant to salinity stress tolerance or avoidance at the vegetative stage. Our overall findings were consistent with the general observation that the cultivated tetraploid *Gossypium* has relatively high tolerance potential to salinity. This was based on generally similar responses across cultivars and landraces to moderate stress level that would otherwise be detrimental to other more sensitive crops and also to the model Arabidopsis. However, screening at higher levels of salinity revealed another layer of variation, suggestive of a hidden potential that should be explored further for use in gene discovery and breeding. An important question that emerged from such hidden potential was the possible contributions of known biochemical and physiological mechanisms that have been uncovered by functional genomics and forward genetics in Arabidopsis and few other crop plants.

This study was conducted with the aims of defining at high resolution the gradient of salinity tolerance potentials across the *core-GDRS* using an integrative physiological, biochemical and whole plant-level phenomics, understanding the critical interactions that may cause either physiological gains or drags, and quantifying the contributions of the major regulators and facilitators of Na^+^ exclusion and transport (*i.e.,* vertical and horizontal) mechanisms, namely *GhHKT1, GhSOS1,* and *GhNHX1*. This study also illuminates the complex but hidden physiological interactions exhibited by the more marginally adapted plant species such as *Gossypium* that may not be revealed by studies using more sensitive crop species and model plants. Lastly, this study establishes the foundation for a network-centered discovery paradigm towards the enhancement of the precision of phenotypic selection in cotton breeding.

## RESULTS

### Salinity tolerance potential relative to genetic diversity

Previous efforts to compare salinity stress responses across different subsets of non-GDRS and GDRS accessions made use of 200mM to 300mM as input concentrations of NaCl in hydroponics (Leidi and Saiz, 1997; Zhang et al., 2014; Peng et al., 2016). Our preliminary studies on a smaller subset of test germplasm revealed that such NaCl levels imposed only mild stress that did not elicit obvious differential reactions across cultivars at the whole-plant level. After one-week exposure, no significant differences across genotypes could be detected based on key growth parameters (Supplemental Fig. S1). However, increasing the NaCl input in hydroponics to 500mM or 750mM using a much wider subset of core-GDRS accessions revealed significant differential responses.

Of the representative germplasm panel, which included twenty five (25) uncharacterized core-GDRS accessions and two (2) known salt-sensitive controls (TX-307 and genome RefSeq TM1), a total of twelve (12) accessions appeared to cover the range of stress tolerance potentials relative to the extent of genetic diversity established previously by SSR-based phylogenetic studies (Fig. 1, A-D; Supplemental Fig. S2; Supplemental Table S1; Hinze et al., 2017). These accessions were chosen to represent the *minimal comparative panel* for all subsequent physiological analyses. From this panel, the accessions SA-0033 (very sensitive/inferior), SA-1055 (sensitive), SA-0881 (moderately tolerant/intermediate), SA-0165 (tolerant), and SA-1766 (very tolerant/superior) were chosen to represent the reference haplotypes for each step in the phenotypic gradient. In the SSR-based allelic diversity plot, the tolerant and sensitive genotypes appeared to have distinct origins, with the genome RefSeq cultivar TM1 being quite distant from the superior core-GDRS accession SA-1766, and from the other reference haplotypes across the phenotypic gradient (Fig. 1E). We hypothesized that the minimal comparative panel including the reference haplotypes are meaningful representations of the various assemblages of positive and negative attributes that may be influencing the observed variation in overall tolerance potentials.

**Figure 1.**
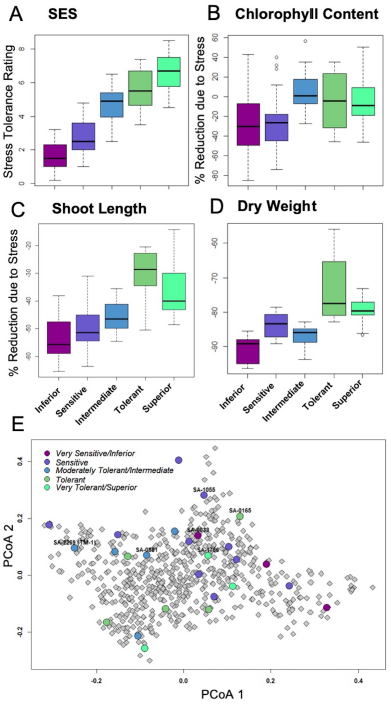
A-D, Boxplots showing significant differences in growth and health parameters with correspondence to the overall stress tolerance potential as represented by the Standard Evaluation Score (SES). Each category across the entire gradient of stress tolerance included at least two genotypes with three to six biological replicates per genotype depending on the assay. E, Principal coordinate plot of the genetic diversity captured by 105 SSR markers across 655 accessions of improved *G. hirsutum* cultivars and landraces (based on Hinze et al., 2016 and Hinze et al., 2017). The twenty-five (25) homozygous accessions of the core-GDRS plus the genome RefSeq TM1 selected for salinity screening are highlighted (circles) with their respective SES as well as other growth and physiological parameters.

### Physiological and biochemical properties contributing to phenotypic gradient

Principal component analysis (PCA) was performed to investigate if the relative stress tolerance ranking across the core-GDRS established by integrating the Standard Evaluation Scoring System (SES) for stress tolerance potential (*i.e.,* 0 to 10 = decreasing severity of stress injury) with various growth parameters could be supported by the inherent variation for other physiological and biochemical properties and their interaction. This analysis was performed using the spatio-temporal profiles for stomatal conductance, membrane injury as revealed by tissue electrolyte leakage index (ELI), lipid peroxidation (LP), tissue Na^+^, K^+^, Mg^2+^, Mn^2+^, Fe^3+^ and Ca^2+^ contents, proline content (Pro), chlorophyll content (Cp), total peroxide content (PRX), total antioxidant capacity (DPPH), and total catalase (CAT) and peroxidase (PER) activities.

Integration of the profiles along the vertical leaf/shoot axis of the plant, *i.e., L1 = oldest/least sensitive, L2 = mid-age/intermediate, L3 = shoot/most sensitive* (Fig. 2), for all physiological, chemical and biochemical parameters showed that at mild to moderate stress imposed by 250mM to 500mM NaCl, three principal components explained 50.4% of the total phenotypic variance (Fig. 3, A and B). The superior and tolerant genotypes (including SA-1766 and SA-0165) were significantly separated from the inferior and sensitive (including SA-0033 and SA-1055) and intermediate (including SA-0881) genotypes along PC1, which explained 25.4% of the total variance. The driving eigenvectors along this axis that also correlated with SES were stomatal conductance, Mg^2+^ and Ca^2+^ content, and total catalase and peroxidase activities, with negative contributions from total peroxide content, and membrane integrity as measured by ELI. The PC2 and PC3 explained 13% and 12% of the total variance, respectively. The PC2 separated the rest of the genotypes from the superior group based mainly on total chlorophyll content, while the PC3 separated the inferior genotypes by virtue of the SES, total peroxidase, and total antioxidant profiles.

**Figure 2.**
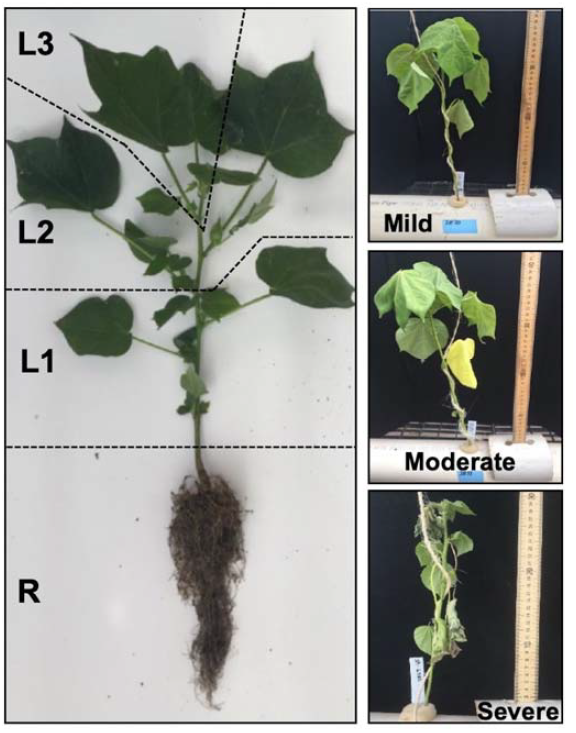
Spatial sampling scheme for the analysis of the vertical distribution of Na^+^ and K^+^, and for profiling the expression of the major genes involved in Na^+^ homeostasis (*GhHKT1*, *GhSOS1*, *GhNHX1*) at mild, moderate, and severe levels of stress. The four sampled positions along the vertical axis of the plant were designated R (Roots), L1 (oldest leaves in the lowest zone of the shoot axis; least sensitive), L2 (mid-age leaves in the middle zone of the shoot axis; moderately sensitive), and L3 (youngest leaves in the uppermost zone of the shoot axis; most sensitive).

**Figure 3.**
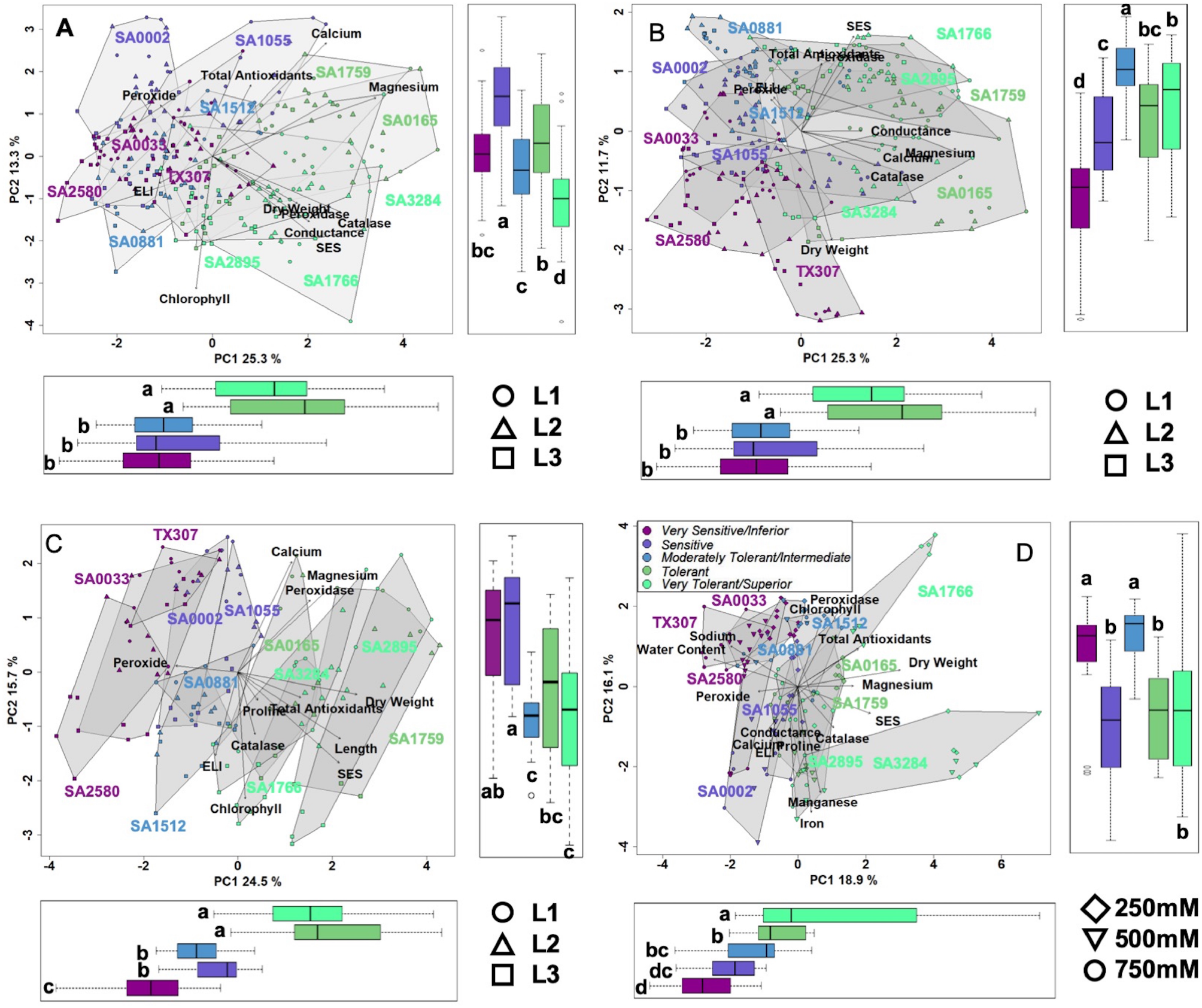
A and B, Scatterplots of shoot PCA at 250-500mM input NaCl. PC1 explained 25.6% of the variation, which significantly differentiates the tolerant and superior genotypes (*R^2^*= 0.505, *p*<0.001). PC2 explained 13.8% of the variation, which significantly differentiates the superior genotypes (*R^2^*= 0.46, *p*<0.001). PC3 explained 12.9% of the variation, which significantly differentiates the inferior genotypes (*R^2^*= 0.40, *p*<0.001). C, Scatterplots of shoot PCA at 750mM input NaCl. PC1 explained 25.4% of the variation, which significantly differentiates the tolerant and superior genotypes (*R^2^*= 0.68, *p*<0.001). PC2 explained 15.9% of the variation, which significantly differentiates the inferior and sensitive genotypes (*R^2^*= 0.33, *p*<0.001), and it also significantly differentiates the leaf positions mostly with L3 (*R^2^*= 0.35, *p*<0.001). D, Scatterplots of root PCA at 250-750mM input NaCl. PC1 explained 18.2% of the variation, which significantly differentiates the genotypes (*R^2^*= 0.50, *p*<0.001). PC2 explained 16.1% of the variation, which significantly differentiates the inferior (*R^2^*= 0.26, *p*<0.001).

At severe stress imposed by input 500mM to 750mM NaCl, the L1-L2-L3 profiles of superior and tolerant genotypes were significantly different from those of the inferior, sensitive, and intermediate genotypes along PC1, which explained 24.4% of the total variance (Fig. 3C). The major attributes driving this separation are SES, relative shoot length, dry weight, total antioxidants, and peroxide content. The PC2 significantly separated the inferior and sensitive genotypes from the intermediate, tolerant, and superior genotypes with 15.8% of the phenotypic variance. Variation within genotype relative organ positions (L1-L2-L3 profiles) was also significantly correlated with this axis.

In the roots (R) under all levels of stress, PC1 significantly separated the tolerant and superior genotypes from the other phenotypic classes, explaining 18.2% of the phenotypic variance (Fig. 3D). The most critical contributors to this axis are SES, tissue Mg^2+^ and Na^+^ contents, dry weight, total peroxide content, and water content. In addition, much of the variation among individuals within the tolerant phenotypic class occurred on this axis and dispersed by the severity of salt stress. The PC2 explained 16.1% of the phenotypic variance and segregated the genotypes belonging to the tolerant classes from the majority of the sensitive genotypes. Tissue Mn^2+^ and Fe^3+^ contents, total peroxidase activity, and total antioxidants are major contributors to this axis.

### Spatio-temporal patterns of Na^+^ and K^+^ accumulation

Root uptake of Na^+^ may occur either through non-selective cation transporters, non-discriminating K^+^ transporters or both. Absorbed Na^+^ can be extruded externally via Na^+^/H^+^ antiporters. The capacity for balancing uptake, extrusion, xylem unloading, and intercellular and intracellular mobilization across less sensitive (L1) to more sensitive organs (L3) are inherent properties whose importance in tolerance have been previously established in other plant species (Fig. 2). To address the potential contribution of these mechanisms to the observed phenotypic gradient across the core-GDRS, Na^+^ accumulation profiles were compared across the minimal comparative panel.

With increasing concentrations of input NaCl in the hydroponics from mild to moderate to harsh levels, the total Na^+^ content in both roots (R) and shoot axis/leaves (L1-L2-L3) also increased (Fig. 4, A-D). The root (R) profiles indicated a slightly higher Na^+^ uptake in the inferior genotypes at all levels of NaCl input than the superior genotypes (Fig. 4A). This trend suggests that superior genotypes may have higher capacities to extrude excessive Na^+^ absorbed by the roots, presumably through mechanisms that may involve Na^+^/H^+^ antiporter systems.

**Figure 4.**
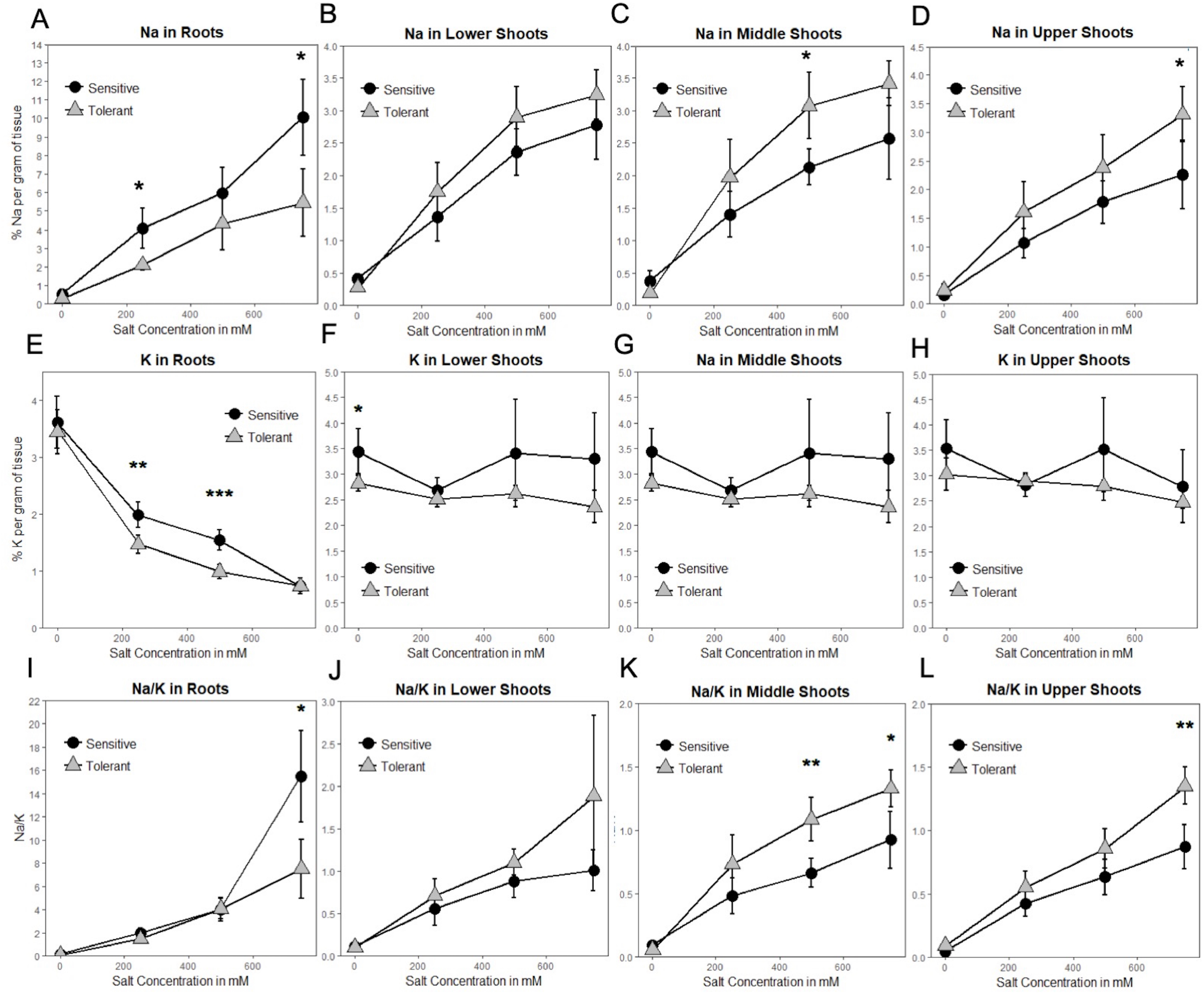
A-D, Line graphs depicting the patterns of Na^+^ accumulation in tissues sampled at increasing input NaCl concentrations along the vertical axis of the plant. Significant increases in Na^+^ concentrations occurred in sensitive plants over time. In the shoots, Na^+^ concentrations increased in the tolerant genotypes. Initially, the Na^+^ increased first in the lower and middle shoots, but as the stress intensified, the sensitive upper shoot accumulated comparable amounts of Na^+^ (_*_ indicates significance at *p* < 0.05). E-H, Line graphs depicting the patterns of K^+^ accumulation in tissues sampled at increasing input NaCl concentrations along the vertical axis of the plant. Significant decreases in K^+^ concentrations occurred in the roots as the stress intensified, which was initially more pronounced in tolerant genotypes. In the shoots, K^+^ concentrations remained constant in all genotypes (significance at _*_*p*< 0.05, \*\**p*<0.01, and \*\*\**p*<0.001). I-L, Line graphs depicting the ratio of Na^+^/K^+^ in tissues sampled at increasing input NaCl concentrations and sampled along the vertical axis of the plant. Significant increases occurred in the Na^+^/K^+^ in roots of sensitive plants over time. In the shoots, Na^+^/K^+^ increased in tolerant genotypes (significance at \**p*< 0.05 and \*\**p* < 0.01).

One peculiar trend observed in the L1-L2-L3 profiles of Na^+^ accumulation was the reversed pattern between inferior and superior genotypes. While this trend appeared to be contradictory to the lower rate of Na^+^ uptake by the roots in superior genotypes, the earlier onset of senescence observed in the older L1 leaves of inferior genotypes may be a contributing mechanism that somehow delays the upward movement and distribution of Na^+^ to the more fragile mid-age L2 leaves (Fig. 4, B, C). However, with progressive increase in input NaCl, the rate of Na^+^ accumulation became more comparable between superior and inferior genotypes. The exception was in the youngest L3 leaves, where the rate of accumulation remained constant in the superior genotypes while tailing off in the inferior genotype (Fig. 4D). Overall, these trends suggest that while root Na^+^ uptake is relatively lower in superior genotypes, the higher Na^+^ accumulation up to the mid-age L2 leaves in superior genotypes particularly indicates that the threshold of sensitivity to Na^+^ toxicity is much lower among inferior genotypes. Thus, lower levels of Na^+^ caused cellular injuries in inferior genotypes while revealing different patterns of spatial Na^+^ accumulation in the superior genotypes.

The profiles of K^+^ accumulation in the L1-L2-L3 axis showed no significant changes across stress levels, genotypes, or organ positions (Fig. 4, F-H). The K^+^ uptake profiles in the roots (R) showed a significant decline with increasing strength of NaCl input, which appeared to be accelerated in superior genotypes (Fig. 4E). Given the flat trend in shoot K^+^ accumulation, increases in Na^+^/K^+^ ratios were essentially determined by Na^+^ accumulation profiles (Fig. 4, I-L).

### Differential expression of *HKT1* genes across the minimal comparative panel

Based on studies in Arabidopsis, one of the important functions of the HKT-type transporters is to facilitate Na^+^ influx to the roots and its recirculation in the phloem (Apse and Blumwald, 2007; Davenport et al., 2007). Given the contrasting profiles of root Na^+^ uptake and shoot axis/leaf Na^+^ accumulation between superior and inferior genotypes (Fig. 4, A-D), the spatio-temporal expression *GhHKT1* was compared across the minimal comparative panel in order to assess their importance to the observed physiological variation.

BLAST queries identified two potential orthologs of the Arabidopsis *AtHKT1* in the tetraploid cultivated *G. hirsutum*, one located on chromosome-1 (A1 in A-subgenome) hence *GhHKT1.A1,* and the other on chromosome-16 (D3 in D-subgenome) hence *GhHKT1.D3*. Each of these orthologs formed monophyletic clades with each of the progenitor diploid *G. raimondii* Ulbr. and *G. arboreum* L. genomes at 99% bootstrap support that corresponded to the appropriate subgenome (Fig. 5A). Together, the *GhHKT1* loci of the three *Gossypium* species were monophyletic with 100% bootstrap support, sister to its closest related taxa in the phylogenetic tree, *Herrania umbratica* (R. E. Schult).

**Figure 5.**
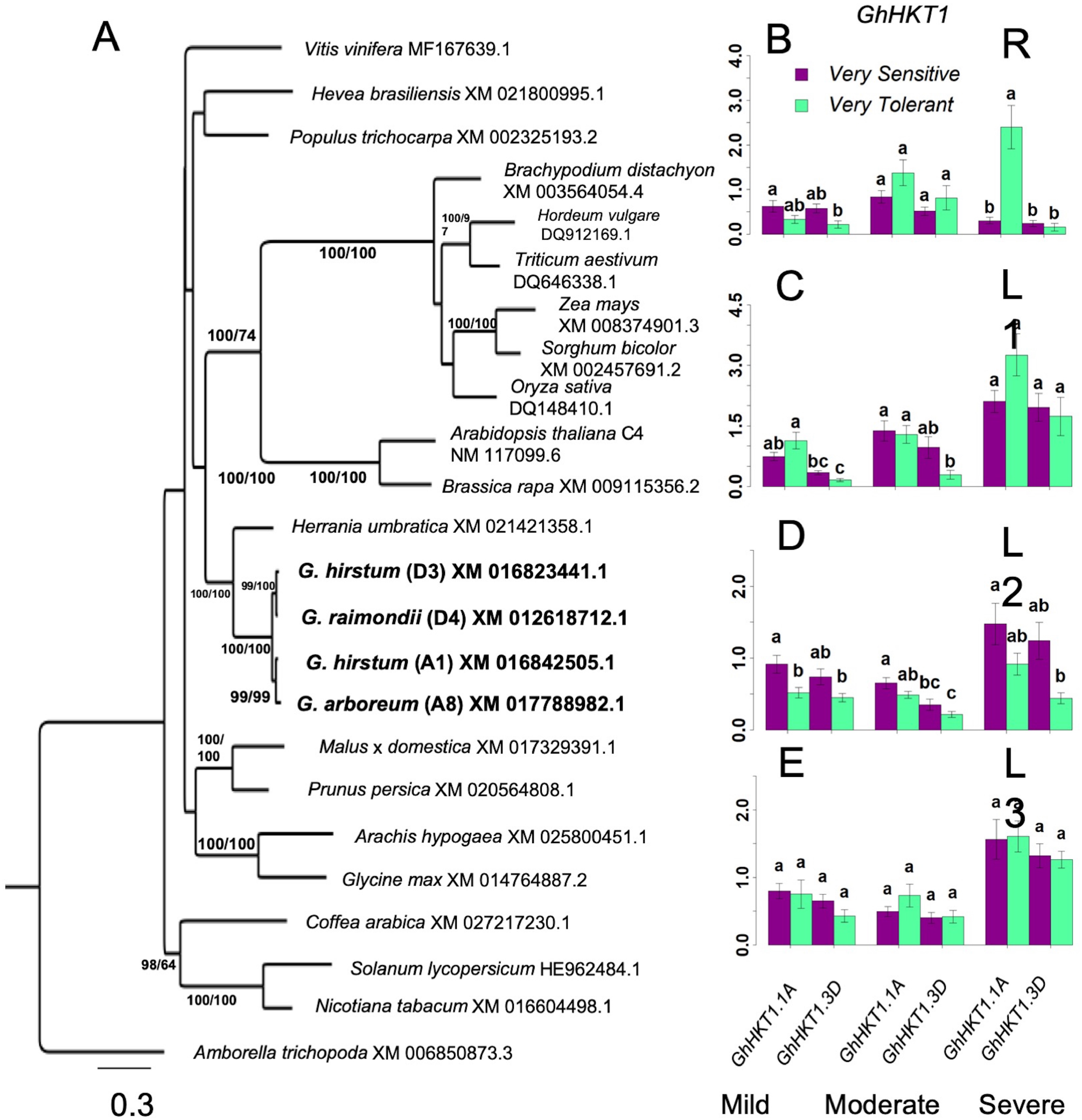
Analysis of *HKT1* homology and its expression in *G. hirsutum*. A, Identification of orthologous *GhHKT1* by maximum parsimony. B, C, D, E, Spatial expression profiles of orthologous *GhHKT1* genes in sensitive and tolerant genotypes along the vertical axis of the plant (B = roots/R; C = leaves/L1; D = leaves/L2; E = shoots/L3). Magnitude of changes in gene expression tend to increase with increasing severity of stress. For the shoots, there are no significant differences in transcript abundances associated with tolerance classification or subgenomic origin of orthologs. In the roots at the highest stress level, there was a significant upregulation of the A-subgenome ortholog (groups assigned by Tukey pairwise comparisons).

Under mild stress, both *GhHKT1.A1* and *GhHKT1.D3* showed slight to moderate downregulation in the roots (R) in both the inferior and superior genotypes, which could be an effect of osmotic shock. With an elevated level of Na^+^ input to impose moderate stress, expression of both *GhHKT1.A1* and *GhHKT1.D3* increased to nearly the control/unstressed levels but still without significant difference between inferior and superior genotypes. Further incremental increase in NaCl input to harsh stress caused significant upregulation of *GhHKT1.A1* only in the superior genotypes, but not *GhHKT1.D3* whose expression remained around the control levels (Fig. 5B). While the superior genotypes appeared to exhibit a unique signature of *GhHKT1.A1* upregulation under high salt, such pattern correlated with increased translocation to the shoots but not Na^+^ accumulation in the roots in the tolerant genotypes, as indicated by tissue Na^+^ content (Fig. 4A) and Na^+^/K^+^ ratio (Fig. 4I). This trend suggests that novel combinations of transport mechanisms could be largely responsible for the observed variation in root Na^+^ uptake across the core-GDRS.

Na^+^ is transported and distributed vertically to the shoot upon absorption through the roots. Analysis of the temporal expression of *GhHKT1* genes across the minimal comparative panel showed that mild to moderate stress levels did not affect the expression of *GhHKT1.A1* nor *GhHKT1.D3* in the leaves, regardless of developmental age and position along the L1-L2-L3 axis in both the inferior and superior genotypes. However, both *GhHKT1* orthologs were significantly upregulated under severe stress in sensitive and tolerant genotypes (Figs. 5, C-E). While HKT-type transporters are known to play some roles in regulating Na^+^ allocation between roots and shoots (Apse and Blumwald, 2007; Davenport et al., 2007), based on the gene expression patterns along the L1-L2-L3 axis and the lack of direct correlation with Na^+^ accumulation profiles (Fig. 4, B-D), the precise contributions of *GhHKT1.A1* and *GhHKT1.D3* to the differential effect of harsh salt stress across inferior and superior cultivars is not clear.

### Differential expression of *SOS1* genes across the minimal comparative panel

The *SOS1* gene is a critical component of the SOS-signaling pathway for regulating cellular Na^+^ homeostasis, encoding a plasma membrane Na^+^/H^+^ antiporter that facilitates Na^+^ efflux (Apse and Blumwald, 2007; Ji et al., 2013). BLAST queries identified three orthologs of the Arabidopsis *AtSOS1* on chromosome-6 (A6 in subgenome-A; *GhSOS1.A6*), chromosome-12 (A12 in subgenome-A; *GhSOS1.A12*), and chromosome-20 (D7 on subgenome-D; *GhSOS1.D7*) of *G. hirsutum* (Fig. 6A). While *GhSOS1.A6* formed a clade with 100% bootstrap support with *G. arboreum*, the *GhSOS1.A12* and *GhSOS1.D7* formed a clade with 99% bootstrap support with *G. raimondii*.

**Figure 6.**
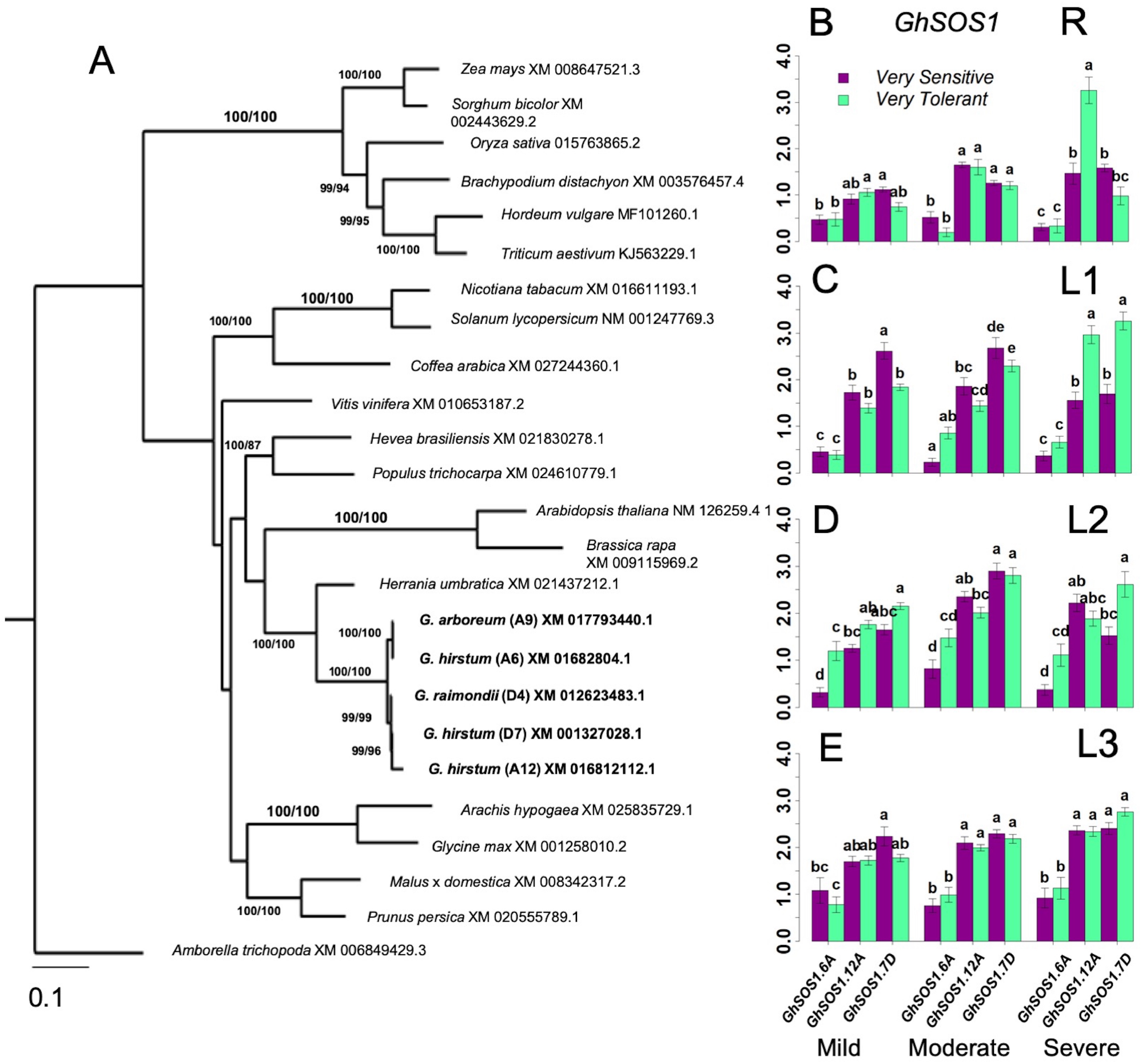
Analysis of *SOS1* homology and its expression in *G. hirsutum*. A, Identification of orthologous *GhSOS1* by maximum parsimony. B, C, D, E, Spatial expression profiles of orthologous *GhSOS1* genes in sensitive and tolerant genotypes along the vertical axis of the plant (B = roots/R; C = leaves/L1; D = leaves/L2; E = shoot/L3). Transcript abundance stayed relatively consistent throughout the stress period along the vertical axis of the plant. The A-subgenome ortholog on chromosome-6 was expressed at half the level of the expression of the orthologs on chromosomes-12 and chromosome-20. Comparable expression of *GhSOS1.A12* and *GhSOS1.D20* initially suggested no expression bias between the subgenomic orthologs. However, according to phylogenies, the *GhSOS1.A12* is most closely related to a gene from *G. raimondii.* Regardless, expression profiles were consistent between tolerant and sensitive genotypes (Groups assigned by Tukey pairwise comparisons).

In the roots (R), expression of all *GhSOS1* orthologs was not affected by mild salt stress in both inferior and superior genotypes. Slight upregulation of *GhSOS1.A12* and *GhSOS1.D7* but not *GhSOS1.A6* were detectable at moderate salt stress (Fig. 6B). Under severe stress, *GhSOS1.A12* exhibited a unique pattern of expression relative to the two other *GhSOS1* orthologs, with significant upregulation specific to the superior genotypes. This positive correlation suggests genotype-specific regulation of *GhSOS1.A12,* and that this gene may have more important contributions to the differential profiles of Na^+^ accumulation in the roots of inferior and superior cultivars, perhaps as a function of the balance between uptake and efflux (Fig. 4A).

Complex patterns of *GhSOS1* expression were observed along the L1-L2-L3 axis. Similar to the patterns in the roots, expression of *GhSOS1.A6* was generally not affected in any level of stress, albeit with slight upregulation relative to the control in both inferior and superior genotypes (Fig. 6, C-E). However, upregulation was detectable for *GhSOS1.A12* and *GhSOS1.D7* under moderate stress, with a tendency for slightly higher magnitude of upregulation in superior genotypes. Interestingly, under severe stress, expression of *GhSOS1.A12* and *GhSOS1.D7* tapered off in older L1 leaves of inferior genotypes but not in superior genotypes where transcript levels continued to rise relative to their levels under moderate stress (Figs. 6C). In younger leaves (L2, L3), different magnitudes of upregulation of both *GhSOS1.A12* and *GhSOS1.D7* were still evident under moderate to severe stress in both inferior and superior genotypes (Fig. 6, D, E). While the temporal and spatial patterns of Na^+^ accumulation in the vertical shoot axis (Fig. 4B) indicated higher levels of accumulation in superior than inferior genotypes across all stress levels, the trends in *GhSOS1.A12* and *GhSOS1.D7* expression tend to suggest that these genes may be involved in some mechanisms that ameliorate the cellular toxicity of excess cytoplasmic Na^+^, perhaps through efflux across the plasma membrane. It also appears that this mechanism is equally functional in both inferior and superior genotypes, albeit at different magnitudes.

### Differential expression of *NHX1* genes across the minimal comparative panel

Sequestration of excessive Na^+^ into the vacuolar compartments through the Na^+^/H^+^ class of endosomal antiporters encoded by the *NHX* gene family has been shown to increase salt tolerance in Arabidopsis through Na^+^/H^+^ and K^+^/H^+^ exchange (Apse et al., 1999). BLAST queries identified three potential orthologs of the Arabidopsis *AtNHX1* in *G. hirsutum*, located on chromosome-2 (A2 in subgenome-A; *GhNHX1.A2*), chromosome-15 (D2 in subgenome-D; *GhNHX1.D2*), and chromosome-17 (D4 in subgenome-D; *GhNHX1.D4*) (Fig. 7A). The *GhNHX1.A2* and *GhNHX1.D2* formed clades with 100% bootstrap support with both ancestral *G. arboreum* and *G. raimondii*. *GhNHX1.D4* only formed a clade with *G. raimondii*.

**Figure 7.**
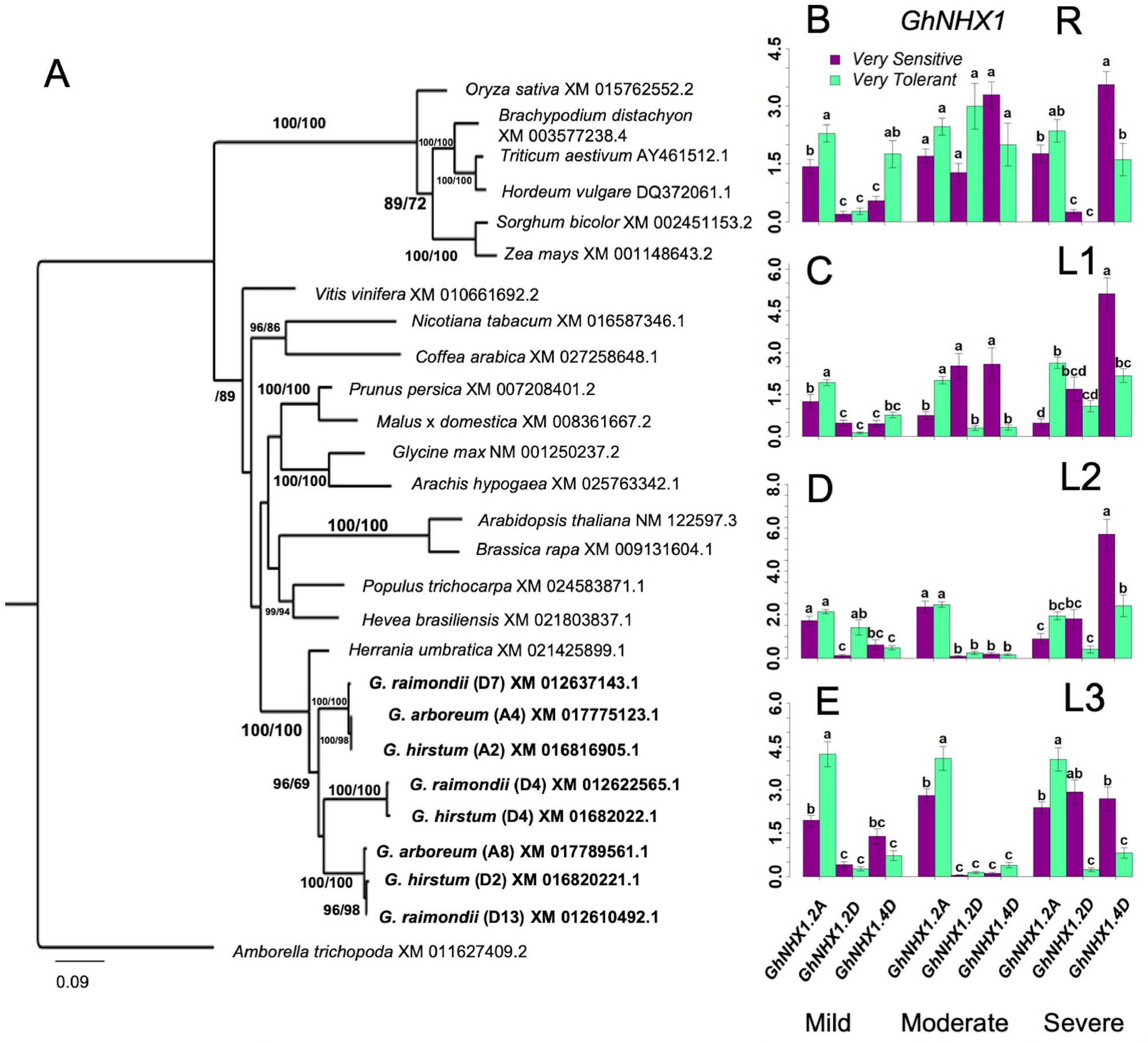
Analysis of *NHX1* homology and its expression in *G. hirsutum*. A, Identification of orthologous *GhNHX1* by maximum parsimony. B, C, D, E, Spatial expression profiles of orthologous *GhSOS1* genes in sensitive and tolerant genotypes along the vertical axis of the plant (B = roots/R; C = leaves/L1; D = leaves/L2; E = leaves/L3). Increases in gene expression was sporadic without overall patterns related to possible variances associated with position along the vertical axis of the plant, stress levels, or tolerance category based on SES. Under severe stress in L1, there the D-subgenome ortholog was upregulated in the sensitive genotypes. There was also a consistent upregulation of the A-subgenome ortholog in the L3 of the tolerant genotype (Groups assigned by Tukey pairwise comparisons).

Compared to *GhHKT1* and *GhSOS1*, the profiles of *GhNHX1* showed greater spatio-temporal variation between inferior and superior genotypes across stress levels. In the roots (R), upregulation of *GhNHX1.A2* and *GhNHX1.D4* but not *GhNHX1.D2* were detected only in superior genotypes under mild stress (Fig. 7B). However, increase in expression became more evident for one or more *GhNHX1* orthologs in both inferior and superior genotypes under moderate and severe levels of stress. Upregulation of *GhNHX1.D2* was also quite evident in superior genotypes only under moderate stress. These trends indicate that while Na^+^ content in the roots vary significantly under all stress levels between inferior and superior genotypes, Na^+^ sequestration to vacuolar compartments appeared to be equally functional in both inferior and superior genotypes. Different *GhNHX1* orthologs also seem to be under distinct regulatory controls in different genetic backgrounds and may be providing some complementation effects.

In the oldest L1 and mid-age L2 leaves, significant increases in the expression of one or more of the three orthologous *GhNHX1* genes were not evident until the plants were subjected to moderate stress, with upregulation occurring in both inferior and superior genotypes for different genes (Fig. 7, C, D). In the youngest L3 leaves, one or more *GhNHX1* orthologs showed significant increases in expression at all stress levels (Fig. 7E). In general, *GhNHX1.A2* had the dominant expression profile under all stress levels and higher in the superior genotypes. While the inferior genotypes had less expression of *GhNHX1.A2*, under the most severe stress, all three *GhNHX1* orthologs were expressed in the more sensitive upper parts of the shoots (L3). Similar to the trends observed in the roots, it appears that different *GhNHX1* orthologs have different patterns of regulation in response to salt stress, and the functions of these orthologs may be complementary across different genotypes. The general trends revealed from the spatio-temporal expression patterns of *GhNHX1* across inferior and superior genotypes suggest that the mechanism of vacuolar sequestration of excessive Na^+^ mediated by NHX1-type pumps are equally functional in both inferior and superior genotypes, hence could not fully explain the observed differential Na^+^ accumulation in the vertical shoot axis of the plant relative to the magnitude of stress sensitivity.

### Na^+^ and K^+^ recirculation capacity of sensitive and tolerant genotypes

In a non-uniform root zone experiment, Kong et al. (2012) previously showed that Na^+^ taken-up by a set of roots in a high-concentration salt solution could be transported by the phloem to a second set of roots connected to the same shoot in a low-concentration salt solution. This transport was blocked by girdling the plants and severing the phloem. The hypothesis that tolerant genotypes are more efficient at Na^+^ recirculation from the shoots back to the roots than the sensitive genotypes was tested by having the inferior and superior genotypes girdled under control and stress (250mM or 500mM input NaCl) conditions. If the tolerant genotypes were more efficient at Na^+^ recirculation, the shoots of its non-girdled plant under stress conditions would have proportionally less Na^+^ relative to its girdled counterpart, and also relative to the sensitive non-girdled and girdled genotypes. Although there were significant differences in Na^+^ concentration between inferior and superior genotypes at different stress levels, differences in the shoot Na^+^ or K^+^ content between girdled and non-girdled plants were not statistically significant (Fig. 8, Supplemental Table S2). In general, the girdled plant declined faster than the non-girdled plants and were not as productive under control conditions. However, this decline could not be attributed to differences in Na^+^ uptake or removal from the shoots. Overall, the results of the girdling experiment implied that superior genotypes may be more efficient in the uptake of Na^+^, but more efficient Na^+^ elimination is not a critical factor in stress avoidance. These results are consistent with the non-significant differences in the expression of *GhHKT1, GhSOS1,* and *GhNHX1* that function as major facilitators of cellular Na^+^ homeostasis (Figs. 5 to 7).

**Figure 8.**
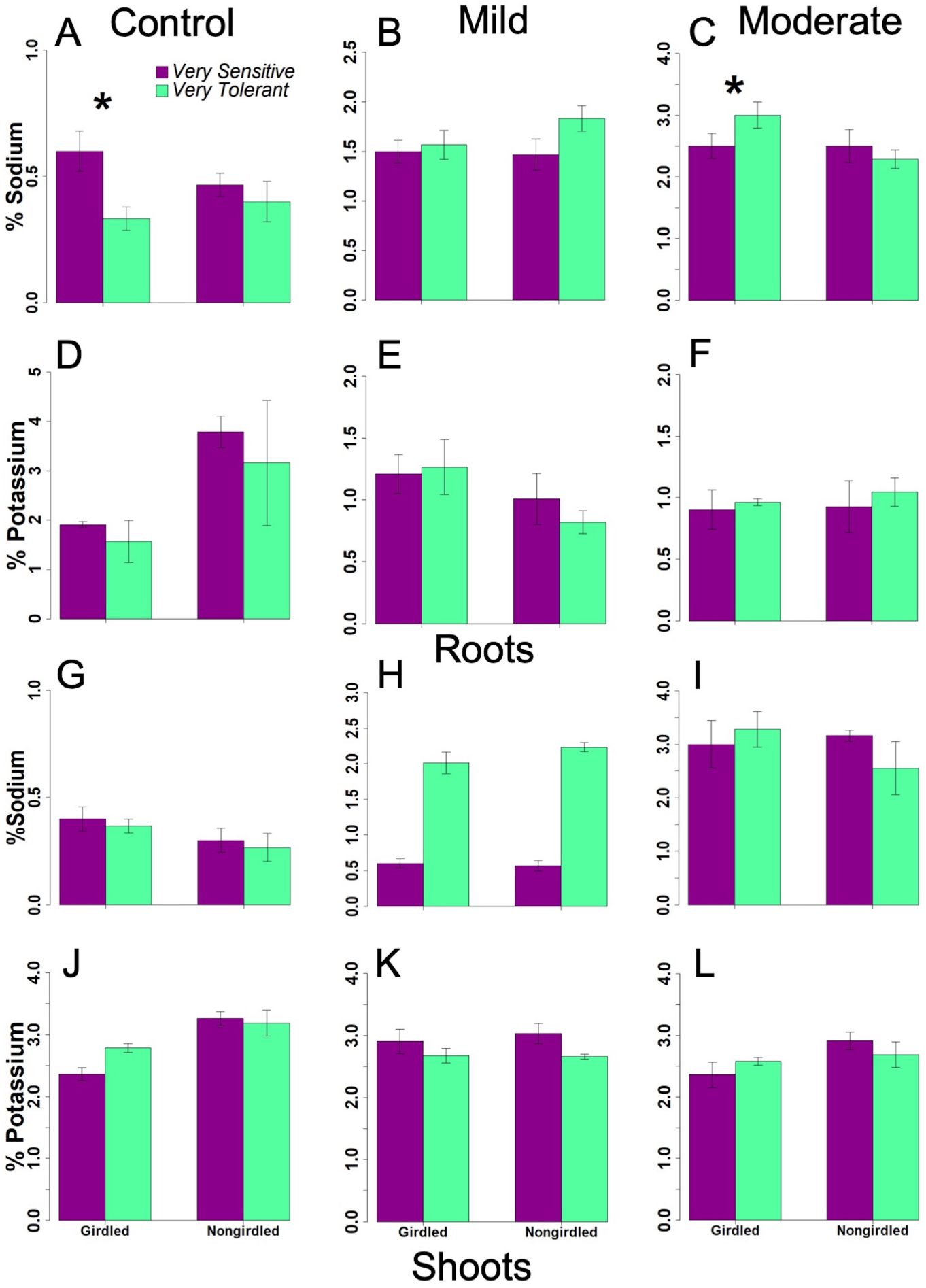
Bar graphs showing the Na^+^ and K^+^ content in the phloem girdled and non-girdled plants under mild and moderate stresses in the roots and shoots. Little to no differences in ion content suggested that recirculation was not a mechanism with significant contribution to improved salinity tolerance potential (Supplemental Table S2).

### Salt tolerance potential explained by multi-factor interaction

The *randomForest* analysis is a machine learning regression tree algorithm that facilitates the classification of different objects into groups by assessing the relative importance of multiple variables contributing to a trait and their interactions through an iterative process (Liaw and Wiener, 2002). Thus, it can be used to weigh the importance of different variables in a set of classifications such SES. This analytical approach was used in this study to integrate all measured physiological, biochemical, and molecular properties across the minimal comparative panel in order to reveal some patterns that mimicked the general trends uncovered earlier by the principal components (Fig. 3; Supplemental Table S3). Among the attributes measured along the L1-L2-L3 axis, variation in total tissue Mg^2+^ content, dry weight, total catalase activity, total peroxidase activity, stomatal conductance, *GhNHX1* differential expression, total proline content, and chlorophyll content had the most significant contributions to the total phenotypic variance across the minimal comparative panel. The contributions of tissue Mg^2+^ content, dry weight and stomatal conductance to the phenotypic variance remained relatively constant as stress increased in severity. The importance of maintaining chlorophyll stability, antioxidant capacity, membrane lipid protection, proline content and *GhNHX1*/*GhSOS1* expression tended to increase with increasing severity of stress, while catalase and peroxidase activities explained the variance more during milder stress and less during severe stress.

Among the various parameters measured in the roots (R) to assess what physiological processes may be contributing more than others to the total phenotypic variance, total peroxidase activity, Na^+^ content, dry weight, total proline content, lipid peroxidation, and *GhHKT1* expression appeared to be the most important as their profiles were generally less affected by the increase in severity of stress. The relative importance of total catalase and peroxidase activities as well as overall antioxidant capacity appeared to decline with increasing severity of stress hence contributed less to the total phenotypic variance.

A theoretical model of the potential interaction collectively driven by the experimental data generated in this study and other information gathered from the literature is summarized in Fig. 9, A, D, G, and Supplemental Table S4. Each square in this model represents a given physiological, biochemical, or molecular attribute, with sizes reflective of their relative contributions to the overall stress tolerance potential (Supplemental Table S3). Interactions between parameters may be positive, negative, or covary, and bold arrows represent significant interactions. The double ended arrows circling back into the square or measured variables represent the latent unmeasured variables and the unexplained variation in the models. Goodness of fit and fit indices for each model are in Supplemental Table S5.

**Figure 9.**
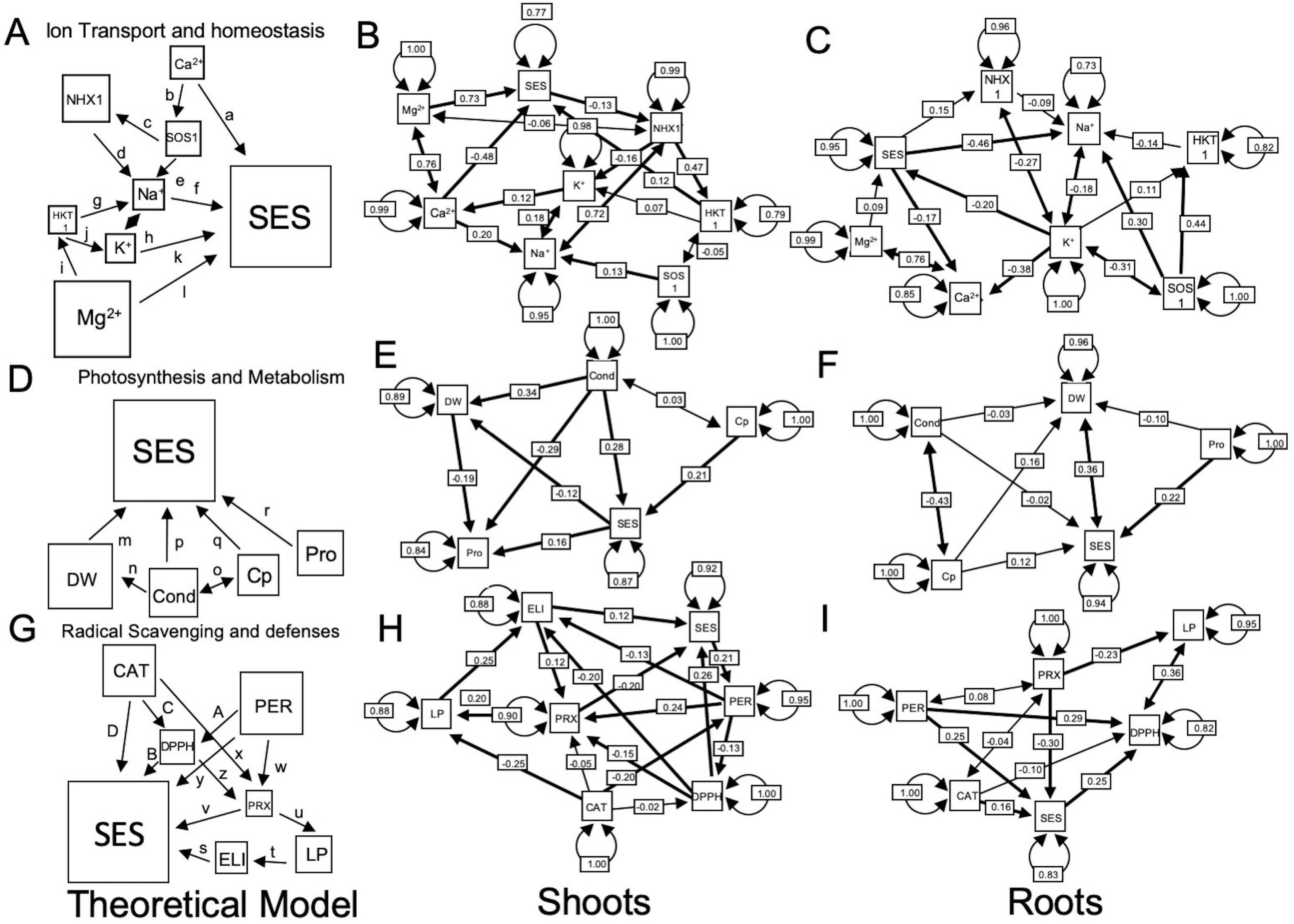
A, D and G, Theoretical models based on referenced studies in Supplemental Table S4. B, C, E, F, H; and I, Empirical models based on data presented.

The general classes or families of physiological outcomes that can be derived from the integration of various variables were *Ion Transport and Homeostasis* (including Na^+^ and K^+^), *Radical Scavenging and Oxidative Defenses*, and *Photosynthesis and Metabolism*. Each respective component contributing to these physiological outcomes are depicted in the model by individual path of coefficients, which also illustrate the multi-factor interactions and how those interactions funneled into the SES as overall measure of stress tolerance potential.

Guided by the theoretical model, a parallel empirical model based on statistically significant interactions uncovered from the dataset was also constructed that incorporates the same general classes or families of physiological outcomes, *Ion Transport and Homeostasis*, *Photosynthesis and Metabolism* and *Radical Scavenging and Oxidative Defenses* (Fig. 9). In the interaction model derived from shoot profiles (Fig. 9B), neither the total Na^+^ content nor the total K^+^ content had a direct impact on SES, while the total Mg^2+^ content and total Ca^2+^ content had direct and significant positive impacts on SES with a strong co-variance between the two variables. The total Ca^2+^ content and *GhSOS1* expression were most correlated with Na^+^ accumulation, which co-varies significantly with total K^+^ content.

Also interesting is the negative correlation of SES with *GhNHX1* expression, and the positive correlation of *GhNHX1* with *GhHKT1* expression, neither of which were suggested by the theoretical model. Moreover, while the empirical model represents a significant improvement over the theoretical model (Supplemental Table S5), at best it only explained 23% of the variance in SES and 5% of the variance in total Na^+^ content. The model for *Ion Transport and Homeostasis* based on the root profiles was closer to the configuration of the theoretical model with the expression of the three Na^+^ homeostasis genes showing correlation with Na^+^ accumulation but only *GhSOS1* was significant (Fig. 9C). Overall, while the statistically significant empirical model better explained the variation in Na^+^ concentration in the root, it still did not explain much of the variation in SES.

In the case of *Photosynthesis and Metabolism*, the empirical model based on the shoot profiles explained only 13% of the variance in SES, mostly by virtue of correlation with stomatal conductance, which explained 11% of the variance in dry weight (Fig. 9E). Stomatal conductance and dry weight also had negative correlations with proline content, while SES had a positive correlation with proline content. Therefore, in the roots, the empirical model was not significantly different from the theoretical model and neither were significant (Fig. 9F).

Based on empirical model for *Radical Scavenging and Oxidative Defenses*, the profiles of antioxidant content and activity and all associated interactions across parameters in the L1-L2-L3 axis was able to explain only as much as 8% of the total variance in SES, with total peroxide content having a negative correlation and total antioxidant content having a positive correlation (Fig. 9H). In the roots, 17% of the variance in SES was explained, but the empirical model was not an improvement over the theoretical model, which explained 23% of the variance (Fig. 9I).

## DISCUSSION

As the understanding of the fundamental mechanisms of salt tolerance continued to advance through functional and comparative genomics in both model and crop plants, the paradigm of a single gene or few genes controlling the complex physiological and biochemical bases of adaptive responses has gradually shifted to more system-wide or holistic paradigm. The modern views suggest that stress tolerance potential of a given genotype is the manifestation of complex interacting biochemical pathways and genetic networks working in concert (Morton et al., 2019). This view is also consistent with the new paradigms proposed by the *Omnigenic Theory* for quantitative traits, which proposed the additive and synergistic effects of several core genes and hundreds if not thousands of peripheral or trans-effect genes (Boyle et al., 2017). Such level of complexity was apparent from the differential responses observed across the cotton germplasm, a plant species known for its inherently high baseline level of tolerance to salinity and other physiologically similar stress effects such as drought. This is evidenced in this study by the high input NaCl concentrations necessary to elicit differential responses and reveal significant variation in sensitivity at the whole-plant level across a meaningful genetic diversity panel such as the core-GDRS.

### Conservation of the first-line-of-defense

While mild to moderate salt stress did not clearly distinguished the sensitive and tolerant genotypes, distinct biochemical and physiological transformations apparently have taken place for either short-term or long-term adaptive mechanisms. For instance, while the total tissue peroxide content begins to build-up in the sensitive genotypes, potentially causing damage to membranes, tolerant genotypes respond by producing more antioxidants, efficient stomatal conductance, and less severe losses in chlorophyll content hence more robust photosynthesis. Interestingly, results of this study also showed a general trend of increased free divalent cations such as Ca^2+^ and Mg^2+^ in the tissues of tolerant genotypes, which has been observed previously among halophytes subjected to NaCl concentrations above 400mM (Vincente et al., 2004). These cations are important components of initial signaling that involves a cascade of protein activation by phosphorylation. For example, intracellular Ca^2+^ plays an important role in the activation of the SOS signaling pathway that controls cellular Na^+^ homeostasis through Ca^2+^-mediated protein phosphorylation (Chinnusamy et al., 2004). Mg^2+^ is an activator of HKT-type transporters such as HKT1 during the facilitation of Na^+^ influx to the roots and its recirculation in the phloem (Chen et al., 2017).

Despite the established knowledge, the general trends revealed in the current study implied that increased ionic concentrations do not necessarily translate to greater magnitudes of expression of the major transport facilitators that regulate cellular Na^+^ homeostasis such as *GhHKT1*, *GhSOS1* and *GhNHX1* in the more tolerant *Gossypium* cultivars. With the exception of minimal correlation of *GhHKT1* with SES in the shoots, there were no other statistically significant or direct relationships revealed by the path analysis between tolerance or sensitivity and expression of the major genes involved in cellular Na^+^ homeostasis. An alternative hypothesis that could address the importance of divalent cations is that they might be contributing largely to osmotic adjustment and reduction of ionic toxicity. Efficient means for maintaining optimal osmotic balance has been implicated with enhanced levels of compatible osmolytes such as proline in tolerant genotypes. While this may appear to be beneficial to the plant as part of the first-line-of-defense or short-term defense, production of these osmolytes often comes with major metabolic expense (Raorane et al., 2015). Our current data suggest that increased production of proline during the early stages of stress or even during mild to harsh stress may translate into major compromises in the overall metabolic status, and ultimately to growth and survival potential of the plant, an effect that appears to be even more pronounced among the sensitive genotypes.

In this study, we have also observed that even the most tolerant genotypes are not necessarily immune to Na^+^ toxicity. The *first-line-of-defense* may delay the onset of injuries, but irreversible injuries may occur at a much later period during chronic exposure to injurious salt concentrations. In the principal component analysis of various physio-morphometric attributes, the profiles of tolerant genotypes under severe stress (750mM input NaCl) tended to overlap with the profiles of sensitive genotypes under mild stress (250mM input NaCl). This implies that sensitive and tolerant genotypes have similar biochemical and physiological responses occurring at different thresholds of osmotic imbalance or ionic toxicity. However, a separate plot for the profiles under 750mM input NaCl showed that the traits that are more closely associated to tolerance such as proline content were still more dominant among tolerant genotypes. Interestingly, while one of the principal components (PC2) segregated the sensitive and tolerant genotypes, separation was much clearer based on the spatial similarities and differences in the L1-L2-L3 axis. As senescence begins in the oldest L1 leaves representing the onset of chlorophyll degradation, such pattern also corresponded to the profile of Ca^2+^ and Mg^2+^ accumulation among tolerant genotypes. At severe salt stress, there is a clear vertical partitioning of resources.

### Cellular Na^+^ homeostasis represents only the baseline component of the untapped potentials in *Gossypium*

The inherent capacity for selective uptake of K^+^ over Na^+^ (HKT1), Na^+^ exclusion (SOS1), and/or avoidance by subcellular sequestration (NHX1) have been viewed as potential sources of genotypic variation for adaptive responses. It is also generally accepted that uptake of Na^+^ by the roots occurs either through passive transport facilitated by non-selective cation transporters, through K^+^ transporters that do not discriminate against Na^+^ when its concentration is high, or through a combination of both mechanisms (Apse and Blumwald, 2007; Davenport et al., 2007; Kronzucker and Britto, 2011). Although there are mechanisms that allow some of the absorbed Na^+^ to be extruded back externally via Na^+^/H^+^ antiporters like SOS1, it has been hypothesized that different plant species and genotypes have inherently unique capacities for controlling the balance between uptake, extrusion, xylem unloading, and intercellular and intracellular mobilization. This balance is presumed to be behind the variation in the net Na^+^ accumulation in sensitive organs such as the leaves and shoots.

Many of the proposed mechanisms for ameliorating the negative impacts of ionic toxicity and osmotic imbalance at the cellular level are based on functional genomic studies in the model Arabidopsis. These studies highlighted the central role of avoidance by maintaining tolerable levels of Na^+^ in the cytoplasm. It has been shown that the more tolerant genotypes tend to accumulate less Na^+^ than sensitive genotypes (Horie et al., 2009; Munns et al., 2012), or that ionic toxicity is mitigated when excess Na^+^ is sequestered in the vacuolar compartment (Aspe et al., 1999; Wu et al., 2004). Furthermore, it has also been proposed that the mechanisms for differential accumulation of Na^+^ could be spatially controlled by retaining much of the Na^+^ absorbed through the roots to the least fragile and relatively more dispensable sink organs of the vertical shoot axis such as the older leaves at the basal region of the plant. This capacity is expected to occur in more tolerant genotypes (Cheeseman, 1988; Munns and Tester, 2008).

Conversely, few other studies have also claimed that the more tolerant genotypes have a general tendency to accumulate more ions under stress, contending that increased ion concentration is necessary to maintain osmotic balance throughout the plant, thereby facilitating new growth. Specifically highlighted in those studies was the maintenance of a lower Na^+^/K^+^ ratio or higher K^+^/Na^+^ (Chen et al., 2007; Shabala et al., 2010). The vertical movements of Na^+^ and K^+^ have been associated with the HKT1-class of ion transporters, which function in ion removal or ion loading from the xylem or into the phloem for recirculation back to roots and eventual extrusion.

In the current study on a meaningful germplasm panel in the cultivated *Gossypium*, spatio-temporal profiles showed that the tolerant genotypes did accumulate more Na^+^ in the shoots compared to sensitive genotypes. This difference was most apparent in the more labile upper/younger organs of the shoot vertical axis (L2-L3) especially at severe stress levels. Accumulation of Na^+^ in the shoots, however, did not vary significantly in the L1-L2-L3 profile, and other than magnitude, there was no significant difference in vertical distribution associated with tolerance. This was consistent with some of the trends in the *GhHKT1* expression in shoot/leaves that showed increased expression with stress level particularly under severe stress (Fig. 5, C-E). As expected, *GhHKT1* expression was highest in L1, but this level of expression did not translate to significant increase in Na^+^ accumulation. There were also no significant difference in *GhHKT1* expression in the shoot/leaves at any stage of stress between the sensitive and tolerant genotypes.

While there was an increase in shoot Na^+^ concentrations, such increases were significantly lower in the roots of the tolerant genotypes compared to sensitive genotypes, and the expression of *GhHKT1.A1* was significantly upregulated in the tolerant genotypes. Considering the multiple functions attributed to HKT-type transporters, the precise implications of these findings are difficult interpret. If Na^+^ was being withdrawn from the xylem sap at a higher rate in tolerant genotypes, then Na^+^ in the roots should increase at a higher rate compared to sensitive genotypes. Na^+^ in the shoot/leaves should increase at a lower rate compared to the sensitive genotypes. Since *GhHKT1.A1* is co-upregulated along with *GhSOS1.A12* during severe stress in the tolerant genotype, it is possible that Na^+^ in the roots was being eliminated from the plant. Had this occurred though, there would have been a reduction in the rate of increase of Na^+^ in the shoots.

K^+^ concentrations remained constant along the L1-L2-L3 axis but declined exponentially in the roots, and the magnitude of such decline at higher stress levels was more pronounced among the sensitive genotypes. Because shoot K^+^ remained constant, the Na^+^/K^+^ ratio increased with increasing Na^+^. The increasing ratio of Na^+^/K^+^ was more pronounced in the roots due to the decreasing K^+^ concentrations coupled with the increase in Na^+^ and being immersed in an environment of ever-increasing Na^+^/K^+^ ratio, *i.e.,* 56/1 at 250mM, 112/1 at 500mM, and 169/1 at 750mM, nearly 10-times higher than the ratios in the sensitive genotypes. Assuming that shoot K^+^ concentrations were stable, the activity of *GhHKT1.A1* in the roots would have been unrelated to K^+^ concentration since it did not change through the course of the experiment. However, if K^+^ leaching did occur in association with transpiration, then *GhHKT1.A1* activity would be necessary to maintain a constant shoot K^+^ concentration and may account for the reduction in the roots.

The downregulation of *GhHKT1* in the shoots at mild to moderate stress levels suggests that Na^+^ recirculation to the roots is not a significant contributor to stress tolerance variation across the minimal comparative panel used in this study. This was confirmed by the phloem girdling experiment, which showed no significant differences in Na^+^ accumulation between girdled and non-girdled plants. The activity of *GhHKT1* however did increase in the shoots at severe stress, and thus Na^+^ recirculation may occur universally late during stress progression, but neither the gene expression nor Na^+^ content data would suggest this has a differential contribution to the observed phenotypic variation in stress tolerance potential.

We were unable to continue the girdled plant experiment into severe stress levels because the girdled plant incurred more stress injuries than the non-girdled plants. However, one observed effect of girdling was that a plant with a severed phloem did not react to the instantaneous effects of osmotic shock. This was typical of all cultivars used in this study, where wilting was apparent within minutes of stress application. The long-distance stress signaling that occurred between the roots and the shoots was terminated with the severed phloem.

When the total Na^+^ accumulation of the roots and shoot/leaves were combined, there were little difference between sensitive and tolerant genotypes, implying that the capacity for exclusion is consistent across the minimal comparative panel. As mentioned above, Na^+^ accumulation was distributed differentially in sensitive compared to tolerant genotypes, with more Na^+^ in the roots and less in the shoots, and accompanied by an increase in *GhSOS1.A12* expression in the roots. *GhSOS1.A12* and *GhSOS1.D7* were also upregulated in the lower L1 leaves under severe stress in the tolerant genotypes compared to the sensitive genotypes. Otherwise, in the upper portions of the shoot axis (L2-L3), there was no significant difference in *SOS1* expression (Fig. 6, D, E). The difference in *GhSOS1.A12* and *GhSOS1.D7* expression may be indicative of early senescence in the lower older shoots of the sensitive genotypes. Overall, *SOS1* activity did increase as the severity of stress increased with no significant implication to reduced Na^+^ accumulation in the shoot that could be meaningfully interpreted in relation to tolerance potential.

Since *GhNHX1* genes function to sequester Na^+^ in the vacuole, their activity is unrelated to Na^+^ uptake but may contribute to total accumulation. In the upper and more sensitive L3 organs of the shoot (Fig. 7E), *GhNHX1.A2* was upregulated compared to all other *GhNHX1* genes in the tolerant genotypes at all stress levels. This may explain why in all tissues, the rate of Na^+^ accumulation declined except in the upper shoots. In the roots through middle organs of the shoot (R-L1-L2), *GhNHX1.D4* was upregulated in the sensitive genotypes during severe stress (Fig. 7, B-D). These tissues at this point in the experiment were severely compromised and such pattern of gene expression may represent a last attempt at defense and survival.

Taken as a whole, Na^+^ transport, mobilization, and sequestration explain little in terms of the overall variation in salinity stress response and tolerance potential across the germplasm. A more reasonable explanation for the increase in Na^+^ accumulation in the shoot was more efficient transpiration under stress. On average, the more tolerant genotypes had 30 mmol/(m²·s) more stomatal conductance than the sensitive genotypes. Overall, there is little evidence to support that exclusion or avoidance play a significant role in differential genotypic responses to salt stress in cultivated *Gossypium*.

### Contributions of the A and D subgenomes

Polyploidy can have a positive effect on abiotic stress tolerance, and this phenomenon has been documented by seminal observations on the hexaploid bread wheat (*Triticum aestivum* L.) and tetraploid durum wheat *Triticum durum* Desf. (Munns et al., 2012). Diploid D-genome species such as *Gossypium davidsonii* Kellogg are known to be salt-tolerant (Zhang et al., 2016). Current theory proposes that domestication of *G. hirsutum* favored the orthologs of the D-subgenome for abiotic stress tolerance mechanisms (Zhang et al., 2015). Comparison of the effects of salt stress on the differential regulation of orthologous genes *GhHKT1*, *GhSOS1*, and *GhNHX1* from the two sub-genomic components of the tetraploid *G. hirsutum* did not show a clear indication that either subgenome exerts a more dominant contribution than the other. Expression of *GhHKT1* genes tend to always favor the A-subgenome orthologs. On the other hand, expression of *GhSOS1* genes tend to favor the D-subgenome orthologs when taking into consideration that *GhSOS1.A12* while located in the A-subgenome originated from the D-subgenome. Additionally, expression of *GhNHX1* genes showed different temporal and spatial patterns of expression, *i.e.,* the D-subgenome orthologs were dominant at severe stress in the inferior genotypes, while the A-subgenome orthologs were dominant in the superior genotypes particularly in the upper L3 organs (Fig. 7E).

Perhaps more intriguing was the relationship of the orthologous genes to their progenitors and their positions in the A and D subgenomes. While *GhHKT1* has two loci corresponding to each subgenome, *GhSOS1* has a duplicate copy of the *G. raimondii* ortholog in *G. hirsutum* on chromosome-12A. *GhNHX1* is another example of the dynamic nature of polyploidy in the sense that *G. raimondii* has three copies, whereas *G. arboreum* only has two. Combining the genomes, we would expect *G. hirsutum* to have five, but instead, there are only three. The *G. raimondii* ortholog that would correspond to the gene copy on chromosome-7 and the *G. arboreum* gene that corresponds to the paralog on chromosome-8 have been lost to maintain the gene copy number similar to the D-genome parent. While cultivated cotton is an allotetraploid, its genome is estimated to be roughly 10% smaller than the combined estimates for the diploid progenitors thus downsizing since the initial tetraploidization event (Wendel et al, 2009). This process has continued through domestication and the germplasm represents genetic isolates that have been under selection for different abiotic stress factors that contributed to the genetic diversity represented by the minimal comparative panel examined in this study.

### Significance of physiological networks to salinity tolerance

At an input NaCl concentration of 500mM and above, we began to see the clear onset of salt stress injury in the oldest L1 leaves of the inferior genotypes first. As the stress continued over time with increasing intensity, the injury progressed through the vertical shoot axis towards L2 and L3. Similar patterns of injury started appearing in the more tolerant genotypes but with significant time delay. It was this observation that led to the central hypothesis that Na^+^ toxicity in the upper portion of the plants and in the more tolerant genotypes was averted by the differential activity of the Na^+^ transport system, which acted to reduce or sequester Na^+^ concentration in the more sensitive organs of the more tolerant genotypes. As the data revealed, genes involved in Na^+^ homeostasis exhibited some patterns of genotype-specific and vertical axis position-specific expression, but these patterns alone did not translate to differential patterns of spatio-temporal Na^+^ accumulation. Instead, the differential responses observed across the minimal comparative panel appeared to be consequences of complex physiological and biochemical interactions.

Of the physiological parameters compared across the comparative panel, total antioxidant capacity, stomatal conductance, chlorophyll content, and divalent cation content explained more of the variance in SES and biomass accumulation than either Na^+^ or Na^+^ transport. Still, within each physiological class, *Ion Transport and Homeostasis*, *Photosynthesis and Metabolism,* or *Radical Scavenging and Oxidative Defenses*, only a small portion of the variance across the minimal comparative panel was explained. Our theoretical model of stress tolerance had SES as an endogenous variable and assumed our measured parameters contributed either directly or indirectly through other attributes acting as components of the total stress tolerance potential. What was unexpected was by reversing the direction of some of these interactions, and by eliminating others, there was a significant improvement in the empirical model relative to the theoretical model. This would suggest that what we have assumed to be attributes that were causal to stress tolerance mechanisms maybe actually be the consequences. For example, higher antioxidant activity did not improve salt tolerance, but instead, healthier plant produced more antioxidants.

The general trends revealed by the *randomForest* analysis indicate that the known mechanisms involved in the *first-line-of-defense* including radical scavenging and Na^+^ efflux and homeostasis contributed to the overall stress tolerance potential across genotypes. However, the relative importance of such mechanisms tended to decline with prolonged exposure to stress and with greater severity of cellular toxicity. Based on these models, there appears to be a much more complex synergy at the cellular, biochemical, and molecular levels beyond the *first-line-of defense* mechanisms that could provide a better quantitative measure of the total phenotypic variance. These types of interactions could not be revealed at their highest possible resolution even by the multi-dimensional datasets.

Much of what we have revealed in this study is the baseline tolerance that makes cotton more tolerant to salt than other crops. Although some of the variations observed in the minimal comparative panel was explained by the multi-dimensional data, the major sources of variation were still not accounted for, indicating that there is an additional layer contributing to the variance yet to be explored. While this study leaves many questions yet to be answered, understanding the foundations for the baseline tolerance is important because it becomes the basis for the interpretation of the more extensive transcriptome analysis which is currently underway. The rapidly emerging paradigm on how to look at complex traits in humans (*i.e.,* Omnigenic Theory) provides a good backbone and inspiration to examine the major physiological and biochemical components (core effects) and the multitude of peripheral or trans-effect components that together account for the totality of the stress tolerance potential of a given genotype (Boyle et al., 2017). This may be accomplished in the future by combining the power of multi-dimensional phenomics with high resolution *expressionQTL* (*eQTL*) and genomic modeling.

## MATERIALS AND METHODS

### Evaluation of salinity stress responses across the core-GDRS

The comparative panel referred to in this study as *core-GDRS* was comprised of twenty-five (25) accessions selected from the US National Cotton Germplasm Collection (College, Station, TX). This germplasm subset encompasses 18 globally distributed locations supported with the total range of allelic diversity and haplotypes across the entire collection as reveled by 105 polymorphic simple sequence repeats (SSR) loci selected using PowerCore (Kim et al., 2007; Hinze et al., 2016). Publicly available SSR datasets were used for Principal Coordinates Analysis (PCoA) to correlate the allelic diversity-based classification with phenotypic variation, *i.e.,* salinity stress tolerance potential (Hinze et al., 2017). The minimal comparative panel represents a further reduced subset selected from the *core-GDRS* based on salinity tolerance ranking and included a known salt-sensitive cultivar TX-307 as baseline control as well as the genome RefSeq genotype TM1 (Li et al., 2015; Zhang et al., 2015).

Salinity stress experiments were conducted under greenhouse conditions at 30-35^°^C/24-26^°^C day/night temperature regime, 20% to 30% relative humidity (RH), and 12h photoperiod with 500 μmol m^−2^s^−1^ average light intensity. Salinity tolerance evaluation was performed using a continuously flowing tube-network hydroponic system customized by Diversity-D Inc. (Brownville, Texas, USA). The hydroponic system has an automated pH and electrical conductivity (EC) measuring features that monitored both parameters according to set conditions at constant intervals during the experiment. Three parallel hydroponic systems with a total capacity of 60 plants each were used, representing two (2) salinity stress and one (1) control experiments. Each experiment was comprised of seven (7) replicate plants that were randomly positioned around the hydroponic tube-networks. The growth medium stock solution was made up of Peter’s Professional Hydroponic Special Fertilizer 5-11-26 (JR Peter’s Inc., Allentown, PA, USA) at full-strength concentration (1 g/L) amended with 0.66g /L CaNO_3_ and adjusted to pH 6.5 with 1mM HCl.

Seedlings were grown in 4-inch deep square seedling pots using standard peat moss potting mix until the fourth node stage (N4). Healthy seedlings were selected and transplanted in the hydroponics system with ¼ strength growth medium and allowed to acclimate until the N5 stage with gradual increase in the concentration of growth medium over a period of one week until full-strength. The first experiment was at moderate stress involving a sequential seven-day application of NaCl to the hydroponics system at an input concentrations of 200mM and 300mM. The second experiment was a harsh stress involving incremental increases of NaCl at an initial input of 250mM, then an increase to 500mM, and then a final input of 750mM with three-day interval between each increment.

### Standard evaluation of salinity stress injury

After five days of severe stress, the high salt solution was replaced with regular full-strength hydroponic solution and plants were allowed to recover for one (1) week. At this time, the overall health status of each plant was assessed qualitatively by assigning *Standard Evaluation Score* (SES) using a scale modified from Gregorio et al. (1997) with decreasing severity of stress injury from 0 to10. SES scoring was performed blindly with each plant referenced according to their positions in the hydroponics experiment matrix to avoid subjective ranking. SES scores were averaged across replicate plants and experiments to generate a mean SES for each genotype.

### Measurement of growth and physiological parameters

Twenty four hours after each incremental increase in input NaCl concentration, the physiological status of each plant was evaluated by measuring multiple parameters. Shoot length (L) was measured from time-0 (*t_0_*) to time-n (*t_n_*) under control and stress conditions, from which the growth rate was calculated with the equation *((L_n_-L_0_)/(t_n_-t_0_)*. Chlorophyll content was measured using a MC-100 Chlorophyll Concentration Meter (Apogee Instruments, Utah, USA) expressed as chlorophyll concentration in units of *μmol m^−2^*. Leaf chlorophyll content was measured twice along the vertical shoot axis at positions L1, L2, and L3 with three (3) replicates per genotype in each of the three parallel hydroponics system (Fig. 2). Stomatal conductance was measured with the SC-1 Leaf Porometer (Meter Group, Inc. Pullman, WA. USA) for a single measurement per leaf with three repeats.

At each time point, five (5) plants per genotype were harvested and fresh weight (FW) was determined separately for roots and shoots. Dry weights (DW) were determined by drying the roots and shoots separately in a 50^°^C incubator for 7 days. FW and DW were used to calculate the water content (%) using the equation *(FW-DW)/FW*100*. Since plant material was limited, water content was used to estimate the biomass content at 250mM and 500mM salt stress.

### Temporal tissue sampling and electrolyte leakage analysis

For all chemical analysis and biochemical assays, plants were sampled at four positions along the vertical axis of the plant, representing different organs (R = roots, L = shoot/leaves) and/or developmental age of shoot organs (L1 = oldest leaves/most insensitive, L2 = mid-age leaves/intermediate, L3 = youngest leaves/shoots/most sensitive) (Fig. 2). For the electrolyte leakage (EL) analysis, 5mm leaf discs were collected at L1, L2, and L3 positions for a total of three (3) plants. The procedures of Ballou et al. (2007) was adopted to determine the % electrolyte leakage from the L1, L2, and L3 leaf discs by measuring the electrical conductivity (EC) in 2 ml of pure nanopure water (16 to18 M∧) before and after stress using a Fisherbrand^TM^ Traceable^TM^ conductivity meter (Thermo Scientific, Waltham, MA, USA). The %EL from intact tissues was determined relative to total tissue electrolytes after boiling, expressed either %EL or electrolyte leakage index (ELI), according to de los Reyes et al. (2012).

### Measurement of tissue peroxide content, and total peroxidase and catalase activities

Total peroxide (PRX) content and total peroxidase (PER) activity was determined using the Amplex^®^ Red Hydrogen Peroxide/Peroxidase Assay Kit (Invitrogen, Carlsbad, CA, USA) with slight modifications from the manufacturer’s protocol (de los Reyes et al., 2012). Briefly, 50mg and 25mg tissues were used for peroxide and peroxidase extraction in 500 *μ*L and 1000 *μ*L of 20mM sodium phosphate (Na_3_PO_4_) buffer at pH 6.5. The modified assay was performed with 300 *μ*L of 0.2 U/mL peroxidase, 120 *μ*L of 100 *μ*M Amplex^®^ Red/DMSO solution, and 18 mL of 1X reaction buffer for the total peroxide determination, and 1500 *μ*L of 2mM H_2_O_2_, 120 *μ*L of 100 *μ*M Amplex^®^ Red/DMSO solution, and 16 mL of 1X reaction buffer for the peroxidase activity. Assays in 50 *μ*L volume were performed in 96-well microplates with absorbance measurements at 560 nm on an iMark™ Microplate Reader (Bio-Rad, Hercules, CA, USA). Total catalase activity was determined using the Amplex^®^ Red Catalase Assay Kit (Invitrogen, Carlsbad, CA, USA) according to manufacturer’s protocols with minor modification. Due to high catalase activities especially on the salt stressed plants, 60 *μ*L H_2_O_2_ was used for the initial reaction with absorbance measurements at 560 nm. Total peroxidase and catalase activities were extrapolated from standard curves with *R^2^ >* 0.97.

### Lipid peroxidation assay

Plant samples were obtained from R, L1, L2, and L3 tissues (100 mg) and extracted with 1.75 mL of 0.1% (w/v) trichloroacetic acid (TCA) by centrifugation at 10,000g at 20°C for 15 minutes. The supernatant (375 *μ*L) was incubated with 750 *μ*L of 20 % (w/v) TCA and 750 *μ*L of 0.5% (w/v) thiobarbituric acid (TBA) for 30 min at 95°C (Jambunathan, 2010). Absorbance of the cooled reaction mixes (200 *μ*L) was determined at 530 nm and 600 nm. The total lipid peroxidation products was calculated using the Beer-Lambert equation *C = A /(ε × l)*, where *A* is the difference in absorbance at 530nm-600nm, ε is the extinction coefficient of 155 mM-1cm^-1^, *C* is lipid peroxide concentration in mM, and *l* is the length of the cuvette in cm.

### Total antioxidant capacity assay

The DPPH (2,2 diphenyl-1-picylhydrazil) method of Prakash et al. (2004) was used to determine the total antioxidant capacity in the R, L1, L2, and L3 tissue samples. Briefly, 100 mg of ground tissues were extracted with 500 *μ*L absolute ethanol. The assay solution contained 1 mg of DPPH per 6 mL of absolute ethanol, combined with 100 *μ*L of the plant extract. Absorbance at 520 nm was determined after 10 min incubation in microplates. For the standard curve, 4 mg/50 mL of L-ascorbic acid was used as the starting solution. The radical scavenging activity was calculated using the formula *[1-(Abs_sample_/Abs_control_)] × 100*.

### Determination of proline content in plant tissues

A procedure modified from Bates et al. (1973) was used to determine proline (PRO) concentration in R, L1, L2, and L3 tissue samples. Briefly, 100 mg of pulverized plant tissues were extracted in 500 *μ*L 3% (w/v) sulfosalicylic acid. The assay solution contained 1.25 g ninhydrin, 80 mL glacial acetic acid, 20 mL 6 M phosphoric acid, and 25 mL 3% (w/v) sulfosalicylic acid. Supernatant from the extraction mixture (100 *μ*L) was combined with 500 *μ*L of assay solution and incubated in 95°C water bath for an hour followed by ice quenching and addition of 1 mL of toluene. After separation of the organic and aqueous phases, absorbance of 100 *μ*L from the organic phase was determined at 520 nm in 96-well microplate. Standard curve started with a 200 *μ*M solution of L-proline.

### Determination of Na^+^ and K^+^ content of plant tissues

A total of five (5) plants for each genotype were sampled at R, L1, L2, and L3. Tissue samples were collected from each plant 24 h after each incremental increase in NaCl input, and then oven-dried at 50°C for 7 days. Dried tissues (1 g) were pulverized and analyzed for Na^+^, K^+^ content as well as nine other elements through the standard nitric-perchloric acid digestion method, measured on an AA unit per Western States Ver 4.00, P-4-20 (AOAC, 1990; A & L Plains Analytical Laboratory, Lubbock, TX).

### Phylogenetic analysis of genes involved in Na^+^ homeostasis

The cDNA sequences for the target genes were imported into Geneious 6.1.6. (Biomatters Ltd., Auckland, New Zealand) and aligned using ClustalW. Maximum likelihood (ML) analysis was performed using the RAxML method through the online CIPRES portal with 1,000 bootstrap replicates (Miller et al. 2010; Stamatakis et al. 2008). Maximum parsimony (MP) analysis was performed with PAUP* 4.0b10, and branch support was assessed with 1,000 non-parametric bootstrap replicates (Swofford, 2001; Felsenstein, 1985). Orthology of gene loci was inferred when sequences were monophyletic within the genus *Gossypium*. Genes from *G. hirsutum* that were monophyletic with *G. arboreum* were inferred to have originated from the A-subgenome, while genes that were monophyletic with *G. raimodii* were inferred to have originated from the D-subgenome.

### RNA extraction and transcript abundance analysis by qRT-PCR

Tissue sampling for the extraction of total RNA were according to the same spatial design used in all chemical analysis and biochemical assays (R-L1-L2-L3). Tissue samples were frozen in liquid nitrogen and total RNA was extracted with the Spectrum Plant Total RNA Kit (Sigma, St. Louis, MO, USA) according to the manufacturer’s protocols. The cDNA synthesis was performed with 1*μ*g of total RNA using the iScript cDNA synthesis kit (Bio-Rad, Hercules, CA, USA).

Gene-specific primers for transcript abundance analysis by qRT-PCR were designed based on *G. hirsutum*, *G. arboreum,* and *G. raimodii* sequences with the closest homology to the Arabidopsis *AtSOS1, AtNHX1*, and *AtHKT1* as well as other eudicot species. Primer-BLAST was used to design specific primers for each homologous *Gossypium* open reading frames which were validated against the annotated *Gossypium* reference genome (Supplementaal Table S6). The qRT-PCR assays for transcript abundance was performed using the SsoFast EvaGreen Supermix (Bio-Rad) in the CFX384 Real-time PCR system with three biological and two technical replicates. Relative gene expression was calculated using the *ΔΔ*Ct method, and normalized using ln(x-1) (Schmittgen and Livak, 2008).

### Severed-phloem and Na^+^ recirculation experiments

The severed-phloem method was used to investigate the amount of Na^+^ recirculation back to the roots from the shoots. This experiment was performed with SA-1766 representing the most tolerant genotype and SA-0033 representing the most sensitive genotype. The experiment was performed by girdling the stem 2 cm above the crown according to the procedure of Kong et al. (2012). Roots and pooled shoots of control, control girdled, stressed, and stressed girdled plants were sampled under 250mM and 500mM input NaCl, and tested for stress effect, girdling effect, and genotypic effect on Na^+^ accumulation. Analysis of the Na^+^ content of tissues was performed using the standard nitric-perchloric acid digestion method (A & L Plains Analytical Laboratory, Lubbock, TX).

### Statistical analysis

All statistical analyses were conducted with R 3.5.2 (R Core Team 2013). Salt tolerance indices were calculated by dividing the individual stress parameters with the corresponding means of the control parameters (Munns, 2002). Individual variables were tested for normality using the Shapiro-Wilks test, and the variables that were highly skewed were transformed to either log or square root scales. Datasets were normalized for univariate (ANOVA) or multivariate analyses. Tukey’s test (Agricolae Package) was used for multiple comparison of means (de Mendiburu, 2019). Multivariate normality was also tested using the MVN package (Korkmaz et al., 2014). Individual parameters were transformed for normal distribution.

Principal component analysis (PCA) was performed using the *prcomp* function to investigate the relationship of multiple physiological, chemical and biochemical properties. Eigenvectors were displayed on the ordination using the *envfit* function in the Vegan Package with a significance cut-off at *p* = 0.05, with the magnitude of the vector indicating significance (Oksanen et al., 2019). The importance of variables was determined using *randomForest* through an iterative machine learning that made use of 1/3 of the data matrix for model training with 1,000 replicates (Liaw & Wiener, 2002). Interaction between variables was investigated to establish a theoretical model for salinity tolerance mechanisms based on the parameters measured and their putative contributions to stress abatement. Models were tested using the structural equation modeling in the *Lavaan Package* in R (Rosseel, 2012). The modindices function was used for model refinement. Models were visualized using *semPlot* (Epskamp, 2015).

## ACKNOWLEDGMENTS

This study was supported by the Texas Tech University-BASF Corporation Project Revolution Program, and the Bayer CropScience Professorial Chair Endowment to BGDR. Germplasm was provided by the USDA-ARS, Crop Germplasm Research Center, College Station, TX.

